# Differentiating Agonists and Competitive Antagonists of the Serotonin Type 3A (5-HT_3A_) Receptor

**DOI:** 10.1101/2023.05.15.540789

**Authors:** Anthony J. Davolio, W.J. Jankowski, Csilla Várnai, B.W.J. Irwin, M.C. Payne, P.-L. Chau

**Affiliations:** Theory of Condensed Matter Group, Department of Physics, University of Cambridge, Cambridge CB3 0HE, U.K.; Centre for Computational Biology, University of Birmingham, Birmingham B15 2TT, U.K.; Institute of Cancer and Genomic Sciences, University of Birmingham, Birmingham B15 2SY, U.K.; Bioinformatique Structurale, Institut Pasteur, CNRS URA 3528, CB3I CNRS USR 3756, 75724 Paris, France

## Abstract

What makes an agonist and a competitive antagonist? In this work, we aim to answer this question by performing parallel tempering Monte Carlo simulations on the serotonin type 3A (5-HT_3A_) receptor. We use linear response theory to predict conformational changes in the 5-HT_3A_ receptor active site after applying weak perturbations to its allosteric binding sites. A covariance tensor is built from conformational sampling of its apo state, and a harmonic approximation allows us to substitute the calculation of ligand-induced forces with the binding site’s displacement vector. We show that it is possible to differentiate between agonists and competitive antagonists for multiple ligands while running computationally expensive calculations only once for the protein.

## 1 Introduction

The serotonin type 3, or 5-hydroxytryptamine type 3, or 5-HT_3_, receptor is a member of the Cys-loop ligand-gated ion channel family which also includes the nicotinic acetylcholine (nACh), glycine, and gamma-aminobutyric acid (GABA_A_) receptors. These proteins are responsible for fast synaptic transmission and are the targets of many neuro-active drugs. The 5-HT_3_ receptor consists of five subunits, arranged around a central ion channel. There are five subunit types, named A to E, though only the A and B subunits have been characterised in detail. The A subunit can be expressed as a homomer, resulting in the 5-HT_3A_ receptor. These receptors are involved in nausea and vomiting caused by radiotherapy and chemotherapy, and competitive 5-HT_3_ antagonists have been used to reduce such vomiting for decades.^1^

The first electron microscopy structure of this receptor appeared in 1995. ^2^ Recently, a number of experimental structures of higher resolution have become available with a variety of ligands bound.^3–6^ In 2018, Basak et al. ^7^ solved the structure of the apo-form of the 5-HT_3A_ receptor to 4.3 Å resolution using cryo-electron microscopy (cryo-EM); no antibodies were bound to the receptor (PDB code: 6BE1). These researchers followed up with a study of the open-channel states of the 5-HT_3A_ receptor by administering 5-HT to the protein (PDB codes: 6DG7 and 6DG8).^8^ Then in 2020, Basak et al. ^9^ solved the structure of this receptor bound to the antagonists alosetron (PDB code: 6W1J), granisetron (PDB code: 6NP0), ondansetron (PDB code: 6W1M), and palonosetron (PDB code: 6W1Y). These structures provide a reference for both the binding site displacement, which serves as an input to our method, as well as the active site changes to compare against after generating our computational predictions.

The dynamics and allostery of the 5-HT_3A_ receptor has been studied in silico. ^10–14^ Trajectories from Yuan et al. ^11^ identified how 5-HT bound to the receptor site and changed the site conformation, and demonstrated that those changes lead to ion channel opening. Guros et al. ^12^ found that 5 mM 5-HT was adequate to activate the receptor. While these studies are invaluable, they are computationally expensive, and would need to be rerun for each ligand to determine its biological effect. In this work, we aim to define the mechanical linkage between binding site perturbation and ion channel opening (gating) using methods which are less computationally demanding. To this end, we have performed a parallel tempering Monte Carlo simulation of the structure 6BE1 using the CRANKITE^15,16^ method and analysed its trajectory to attempt to define agonists and competitive antagonists and their associated direction of conformational change.

## 2 Methods

### 2.1 Structure

The starting structure of the simulation is the cryo-EM structure of the apo-form of the 5-HT_3A_ receptor, PDB code 6BE1.^7^ We choose this structure because its has been solved without any extra proteins attached to it and, furthermore, the same research group has also solved the structure of this receptor in the open-channel state and when it is bound to four different kinds of competitive antagonists. This structure is shown in Figure 1.

**Figure 1.**
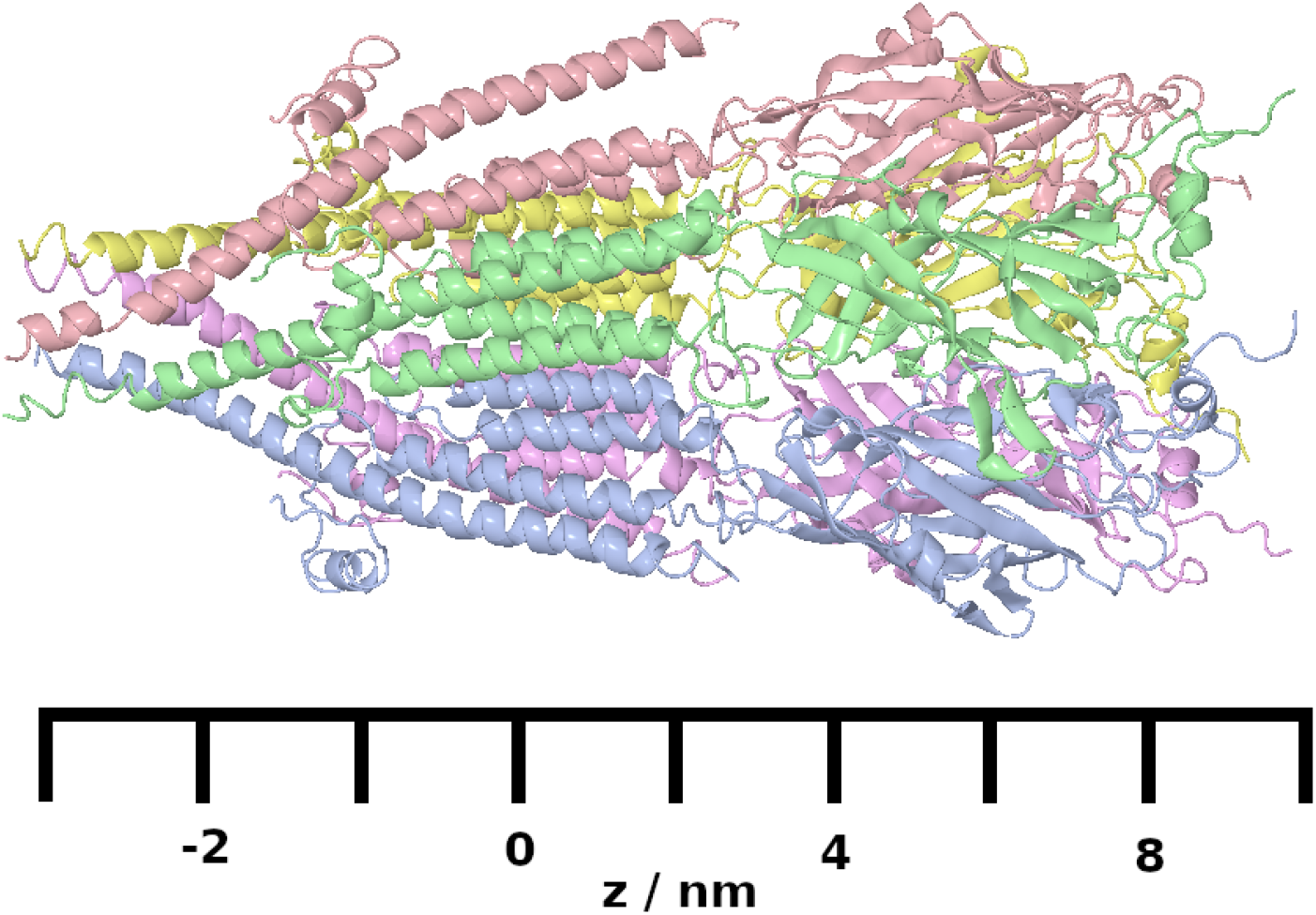
Diagram showing the 5-HT_3A_ receptor with a coordinate system aligned along the receptor’s long axis. The extracellular domain is in the region *z >* 25 Å, the transmembrane domain is in the region −25 Å *z*− 25 Å and the intracellular domain is in the region *z <* 25 Å. The agonist and competitive antagonist binding site is situated between individual subunits and is in the region *z* = 50 Å. These *z*-values will be referred to later in the paper when describing different parts of the receptor.

For comparison with open-channel states, we use the structures of the 5-HT_3A_ receptor bound to 5-HT.^8^ For structures bound to antagonists, we use the structures of the 5-HT_3A_ receptor bound to alosetron (PDB code: 6W1J), granisetron (PDB code: 6NP0), ondansetron (PDB code: 6W1M), and palonosetron (PDB code: 6W1Y).^9^

### 2.2 Simulations

For the molecular simulations, we used CRANKITE,^15^ a Monte Carlo simulation programme that employs a Go-like coarse-grained force field. We chose CRANKITE, because its Monte Carlo move set consists of local moves of crankshaft rotations of the protein backbone and changes of the side chain dihedral angles, the former of which allows a fast conformational sampling of proteins. CRANKITE uses a full atom representation of the protein backbone, together with explicit side chain *β* atoms and *γ* atoms, to include entropic contributions arising from the torsional flexibility of side chains. Each amino acid is treated as hydrophobic or amphipathic. The *γ* atoms represent the rest of the side chain, with an elongated distance between *β* and *γ* atoms to place them near the centre-of-mass of the side chain. In the energy function, the volume exclusion of beads, hydrogen bonds between backbone amide H atoms and carbonyl O atoms, and hydrophobic interactions between *γ* atoms are modelled explicitly. The secondary structure of the protein is held together by an additional energy term, which keeps the backbone of *α*-helices and *β*-strands at the correct backbone twist, and a Go-like contact potential term keeps *β*-sheets intact. The secondary structure of the protein is predetermined and does not change within the simulation.

We used the force field parameters optimised previously,^16^ with the addition of a conical external potential on the transmembrane and intracellular regions to mimic the membrane, the force field^17^ had been optimised using a set of non-transmembrane proteins. The energy contribution of this external potential on an amino acid in the transmembrane and intracellular regions region is:

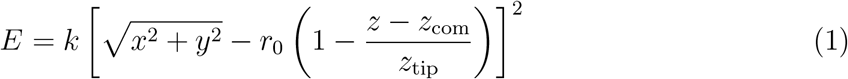

if *z/z*_tip_ ∈ [0, 1] and 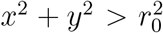, otherwise *E* = 0, where *x, y*, and *z* are the coordinates of the centre-of-mass of the amide N, C_*α*_ and carbonyl C of the amino acid, *k* = 100*RT, r*_0_ = 45 Å, *z*_tip_ = −120 Å, *z*_com_ is the *z*-coordinate of the centre-of-mass of the whole protein where the pore of the protein was aligned to the *z* axis with the intracellular domain pointing in the positive direction as illustrated in Figure 1. The shape of the potential was chosen such that it helps maintain the initial structure of the 5-HT_3A_ receptor, but it does not squeeze the pore closed (there is a minimal energy penalty with the receptor in the initial state).

To explore the possible conformations of the protein while keeping its secondary structure intact, we used parallel tempering simulations with 32 temperature levels ranging from 310 K to 1 023 K, sampling conformations at the lowest temperature level. The simulation was performed for 10^6^ steps. After the simulation finished, we carried out analysis to confirm that the system was equilibrated after 220 000 steps. To allow us a safety margin, useful data were collected from step 240 000. We then performed data analysis of the trajectory using the following methods:

### 2.3 Correlation Tensor

To evaluate how the movements of one amino acid correlate with the movements of another, we construct a correlation tensor, as in Várnai et al.. ^17^ We coarse-grain the CRANKITE trajectory to individual amino acids, whose position becomes its center of mass. We assume that each amino acid has a set of principal directions which can be defined by an ellipsoidal cloud of coordinates **x**_*at*_ = [*x*_*a*1*t*_, *x*_*a*2*t*_, *x*_*a*3*t*_] for the *a*^th^ amino acid, *t*^th^ snapshot, and 1^st^, 2^nd^, or 3^rd^ dimension. These orthonormalized principal axes of motion vectors are equivalent to the eigenvectors of the self-correlation matrix for amino acid *a* in cartesian [*x, y, z*] coordinates. Note that these are spatial correlations rather than temporal correlations. We construct a 3 × 3 correlation matrix for each pair of amino acids *a* and *b*, denoted as 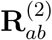 whose elements are given by

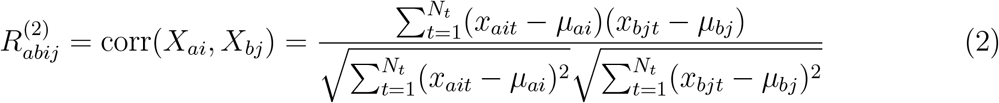

with *i* and *j* as the *i*^th^ and *j*^th^ principle axes of motion of amino acids *a* and *b* respectively, and *μ* as the average position across all *N*_*t*_ snapshots.

### 2.4 Impulse Response

#### 2.4.1 Time-Independent Linear Response Theory

Since our simulation is not consecutive in time, with all computed expectation values corresponding to ensemble averages, we use a time-independent linear response theory developed by Ikeguchi et al. ^18^ to predict the protein’s response to ligand binding. Physically, time-averages of protein structural configuration should coincide with corresponding ensemble averages as a result of ergodicity, which we expect to hold in classical systems such as proteins. For weak protein-ligand interactions, we can approximate our response to first order in the forces

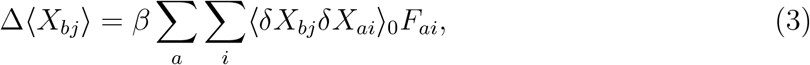

where Δ⟨*X*_*bj*_⟩ is the expected displacement of amino acid *b* in direction *j* induced on binding the ligand, *β* = 1*/k*_*B*_*T*, ⟨*δX*_*bj*_*δX*_*ai*_⟩_0_ is the covariance between *X*_*bj*_ and *X*_*ai*_ at equilibrium (mean position) with no ligand, which multiplied by *β* is a mechanical susceptibility for the response in amino acid positions due to the external ligand forces acting on a protein, consistent with the fluctuation-dissipation relations. *F*_*ai*_ represents the ligand-induced force on amino acid *a* in direction *i*. In the case of strong interactions, we expect an analogous response, captured by a proportionality relation, to hold

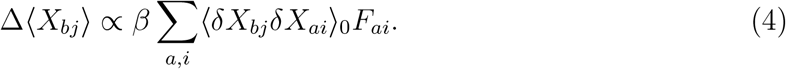

The relation follows from expanding the displacements to higher order in the forces, obtained from a Taylor expansion of the generating functional *Ƶ*(*β*, {*F*_*ai*_}) in terms of forces around zero force limit *F*_*ai*_ = 0.

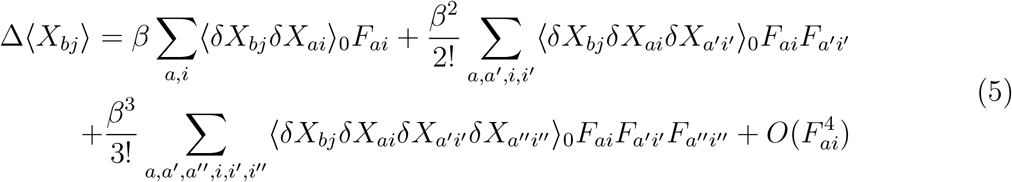

The generating functional is obtained by adding a coupling term *V*_coupling_ = − Σ_*a,i*_ *F*_*ai*_Δ*X*_*ai*_ to the protein Hamiltonian ℋ_0_, which is quadratic in the amino acid displacements within harmonic approximation, exponentiated in a partition function of the ligand-free protein 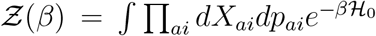, where *p*_*ai*_ denote components of amino acid momentum in units of the Planck constant. The quadraticity of ℋ_0_ allows us to apply Wick’s probability (Isserlis’) theorem, which holds for Gaussian variables, by which all terms odd in fluctuations (e.g., ⟨*δX*_*bj*_*δX*_*ai*_*δX*_*a′i′*_ ⟩_0_) vanish. At the same time, any even term can be expanded in products of all possible permutations of two-amino acid displacement correlators. In the case of the fourth moment (cokurtosis) for example,

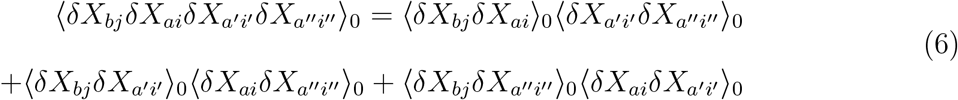

Having applied Wick’s theorem, the higher moments can be collected into a proportionality factor 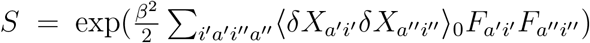, as suggested by Punia and Goel ^19^. Hence, explicitly, strong interaction amounts to

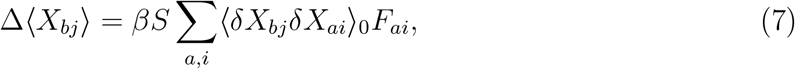

where in the limit of small forces *F*_*ai*_ → 0, by inspection *S* ≈ 1, restoring the weak interaction limit.

#### 2.4.2 Linear Force Approximation

The potential *V*_int_ which describes the amino acid interactions in the ligand-free protein can be Taylor expanded around the equilibrium positions to

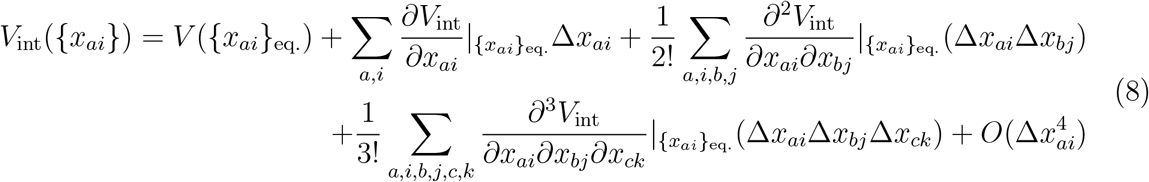

where {*x*_*ai*_}_eq._ corresponds to the average equilibrium positions of amino acids in the protein without a ligand, {*x*_*ai*_}_eq._ ≡ {*x*_*ai*_}_0_, and Δ*x*_*ai*_ denotes a displacement of amino acid *a* in direction *i*. The first term is a trivial constant, the second one vanishes by the equilibrium condition, i.e., derivatives of the potential vanishing at its minimum. In the harmonic approximation, we disregard the terms of higher than quadratic order, as these represent anharmonicity. We define spring constants

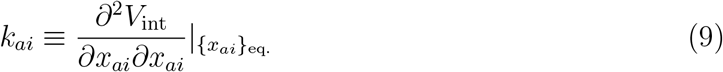

as well as interaction forces on amino acids due to the collective effects of all other amino acids,

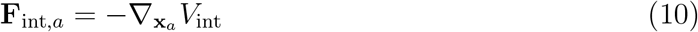

component-wise yielding 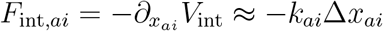.

We assume that, upon inserting the ligand, the new configuration satisfies *F*_*ai*_ = −*F*_int,*ai*_ within the binding site, where *F*_*ai*_ represents a ligand-induced force due to the interaction of amino acid *a* with the ligand. Hence, utilising the harmonic approximation, we obtain *F*_*ai*_ = *k*_*ai*_Δ*x*_*ai*_ for amino acids *a* ∈ the binding site. In this context, {Δ*x*_*ai*_} denote displacements of these amino acids from their apo-state equilibrium positions.

Thus, the ligand-induced forces can be approximated as long as we know the spring constants *k*_*ai*_ and the displacement Δ*x*_*ai*_ of the amino acids in the binding site (either from experiment or docking). CRANKITE simulations provide us with the variance 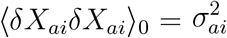, and we can apply the equipartition theorem in the protein described with the Gaussian model, given *V*_int_ consists only of quadratic degrees of freedom to give

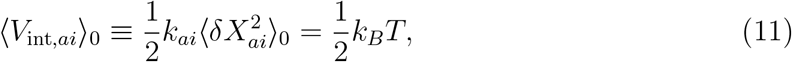

for each component of amino acid displacement **X**_*a*_giving 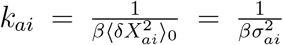. Having deduced spring constants from the ensemble of protein structures, as well as displacements around the binding site Δ*x*_*ai*_, we can infer approximate ligand-induced forces *F*_*ai*_, and therefore other amino acid displacements not in the binding site using the linear response theory introduced in the section above. However, we emphasise that in this harmonic approximation, we neglect entropic contributions to the force, which close to the positionally constrained binding centre, we assume to be negligible.

We can further justify the validity of the harmonic potential and linear force approximations. The statistical relationship between the covariance and the correlation coefficient of *X*_*bj*_ and *X*_*ai*_ is given by

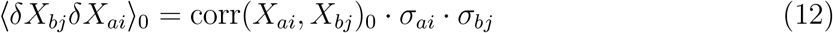

where *σ*_*ai*_ represents the standard deviation of the position of amino acid *a* in direction *i*. Using the above harmonic approximation *F*_*ai*_ = *k*_*ai*_Δ*x*_*ai*_ gives

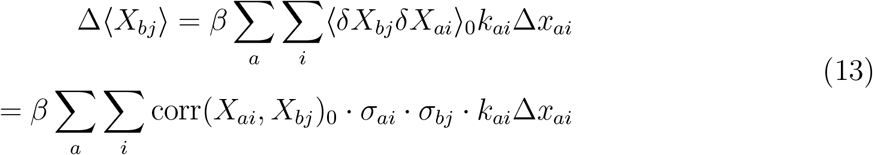

Where the set of {*X*_*ai*_} represents the positions of amino acids across the entire protein and {*x*_*ai*_} represent the positions of amino acids in the binding site, taken to be any amino acid within 6 Å of the ligand. Let us explore a small induced movement of only a single amino acid *a* along only one of its principal axes of motion *i*, represented as Δ*x*_*ai*_, we wish to find Δ⟨*X*_*bj*_⟩ in the special case that *b* = *a* and *j* = *i*. Since self-correlation coefficients are equal to one and we can remove the summation because no other amino acids are externally perturbed under the assumption, we arrive at

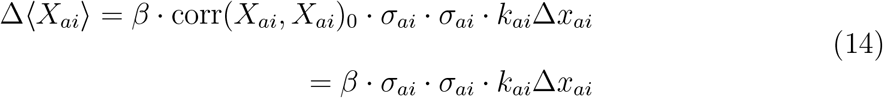

The predicted reactive displacement would be equal to the actual displacement caused by ligand-induced forces in this case, Δ⟨*X*_*ai*_⟩ = Δ*x*_*ai*_. Solving for *k*_*ai*_ we obtain

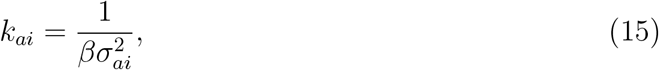

which is equivalent to the previous equipartition theorem result within the harmonic approximation. Finally, reinserting the previous equation into the time-independent linear response relation gives

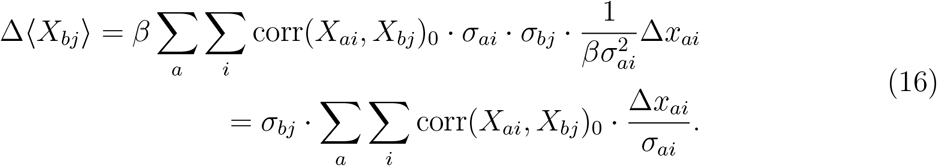

The result is highly intuitive in that one can normalize the input Δ*x*_*ai*_ by its standard deviation *σ*_*ai*_, apply its correlation coefficient with *X*_*bj*_, then analogously normalize the output by its standard deviation *σ*_*bj*_ to obtain an induced response Δ⟨*X*_*bj*_⟩. Overall, this framework allows us to reconstruct the displacements of amino acids not in the binding sites from the displacements of amino acids in the binding site. To perform the approximations required by linear response theory, we have to assume that the ligand induces only a weak perturbation. Therefore, we do not account for major structural rearrangements. However, these are not present between the apo- (PDB code: 6BE1) and holo- (PDB code: 6DG8) form of the 5-HT_3*A*_ receptor.

#### 2.4.3 Response Measurement

We estimated the pore radius of the receptor at a set of sampling points along the *z*-axis (the pore axis) to which the protein ion channel was aligned as illustrated in Figure 1. Samples were taken at 1 Å intervals from −80 Å *< z <* +100 Å; this range covered the main parts of interest of the protein. At each sampling point the ten nearest-neighbour residues were considered and the approximate exponentially smoothed radius was reported as:

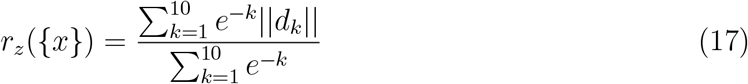

where *d*_*k*_ is the distance from the pore axis to the *k*^th^ nearest neighbour.

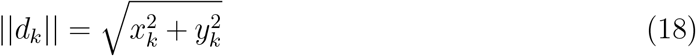

The movements of these amino acids are non-linear and it is quite possible that the M2 helices rotate upon activation with respect to an axis not coaxial with the protein’s pore axis. We note that our choice of a cartesian coordinate system results in any magnitude of forces causing some amino acids to move in towards the channel and some to move out. Thus we must keep the selection of amino acids *k*, and their respective weights *e*^−*k*^, consistent between the apo- and holo-form of the protein, else we risk improperly favoring the amino acids that move inwards in our radius calculations.

### 3 Results

Analysis using auto-correlation functions showed that the CRANKITE equilibration period finished at 220 000 steps of the 10^6^-step simulation. We used the last 760 000 configurations to construct a covariance tensor for the displacement of every amino acid along its principal axes of motion. We used the displacement from the experimental apo-binding site (PDB code: 6BE1) to the experimental holo-binding site to approximate ligand-induced forces and predict the protein’s response to ligand binding for the agonist serotonin (PDB code: 6DG8), and four antagonists – alosetron (PDB code: 6W1J), granisetron (PDB code: 6NP0), ondansetron (PDB code: 6W1M), and palonosetron (PDB code: 6W1Y).

### 3.1 Agonist binding

For comparison purposes, we calculated the channel radius profile for the experimental apo-structure 6BE1; and compared it with the experimental holo-structure 6DG8; this is considered the experimental baseline. Figure 2 shows that binding of five 5-HT molecules to the 5-HT_3A_ receptor caused the ion channel to increase in radius, especially in the transmembrane domain and the extracellular domain. The increase in the transmembrane region radius was about 1.5 Å. We notice a large increase in channel radius around *z* = −20 Å. There are reductions in the channel radius around *z* = −40 Å, *z* = −25 Å, *z* = 35 Å and *z* = 65 Å.

**Figure 2.**
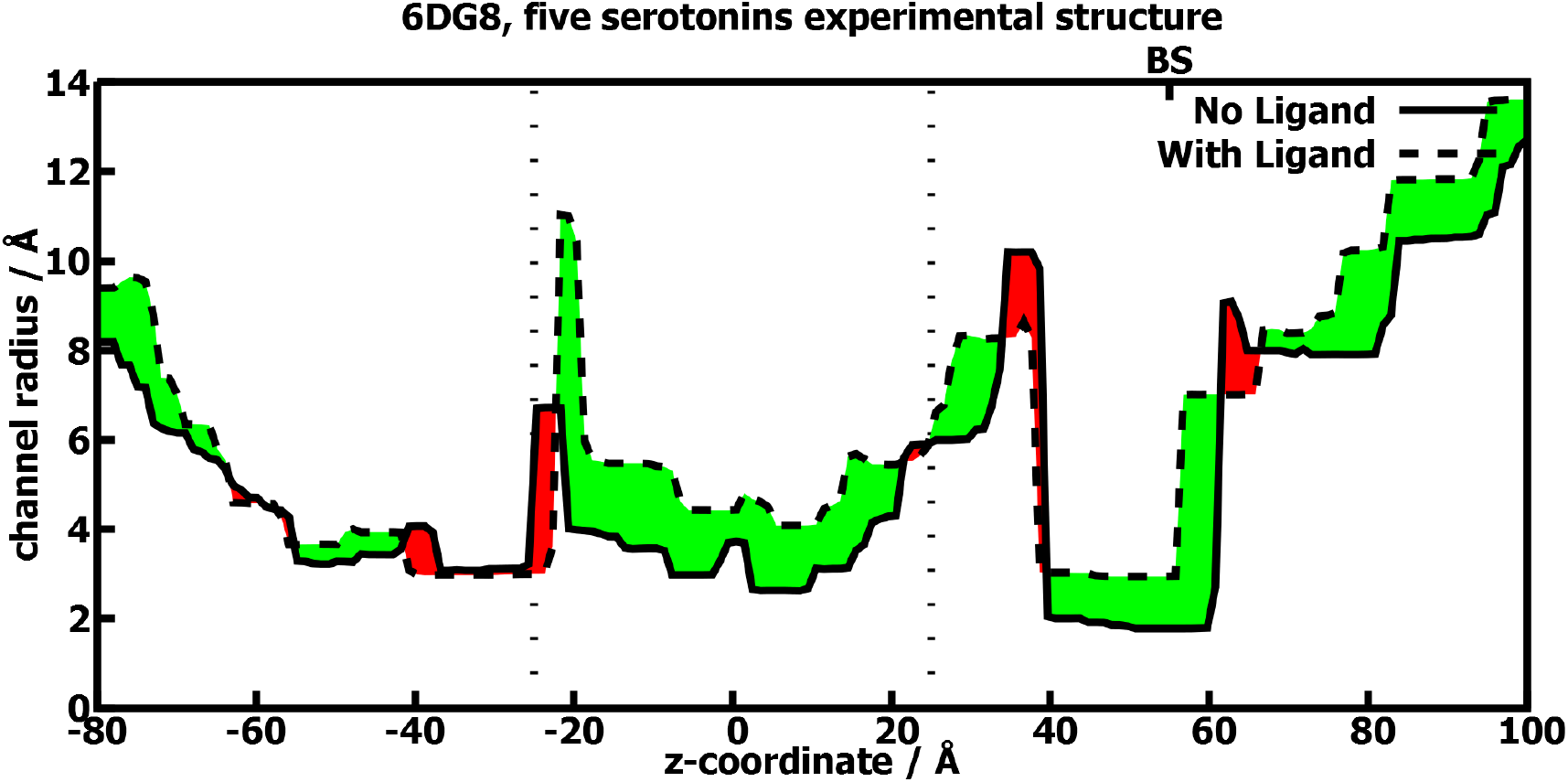
Diagram showing the radius of the ion channel along the length of the 5-HT_3A_ receptor. The solid line represents the data for the experimental apo-structure 6BE1. The broken line represents the profile for the receptor bound to five 5-HT molecules 6DG8. The transmembrane domain, in the region − 25 Å*< z <* 25 Å, is denoted by the broken vertical lines. BS denotes the position of the binding site. The spaces between the solid and broken lines are filled with red (reduction of ion channel radius) or green (increase of ion channel radius).

We then evaluated the channel radii of the predicted structures along their long axis. We used the equilibrium apo-structure, *μ*, averaged from the 760 000 configurations from the data-collection period as the baseline structure. We then used our method to predict the protein structure for one 5-HT to five 5-HT molecules bound (see Supplementary Figures).

We observe that applying 5-HT forward vectors to different numbers of binding sites caused the ion channel to open, especially in the transmembrane domain. We observe that two serotonins provide more activation than one serotonin, and that applying forward vectors to two adjacent binding sites triggers a stronger response than when applied to non-adjacent binding sites. Since the experimental structure 6DG8 has five serotonins bound, we should compare the experimental results with five 5-HT bound (Figure 2) and simulation results with five 5-HT bound (Figure 3a). The only common point is that the ligands caused an increase of the pore width in the transmembrane domain, but in other domains, the simulation results are different from the experimental results. Interestingly, the experimental holo-structure (Figure 2) is most similar to the simulation results for three 5-HT bound in adjacent sites (Figure 3b): both showed a general increase in the ion channel radius of the transmembrane domain up to *z* = 30 Å, an increase in the extracellular domain except where *z* = 35 Å, and a decrease in the region −40 Å *< z <* −25 Å. We also note that previous electrophysiology experiments showed that three 5-HT molecules were required to achieve maximal gating efficacy. ^20–22^

**Figure 3.**
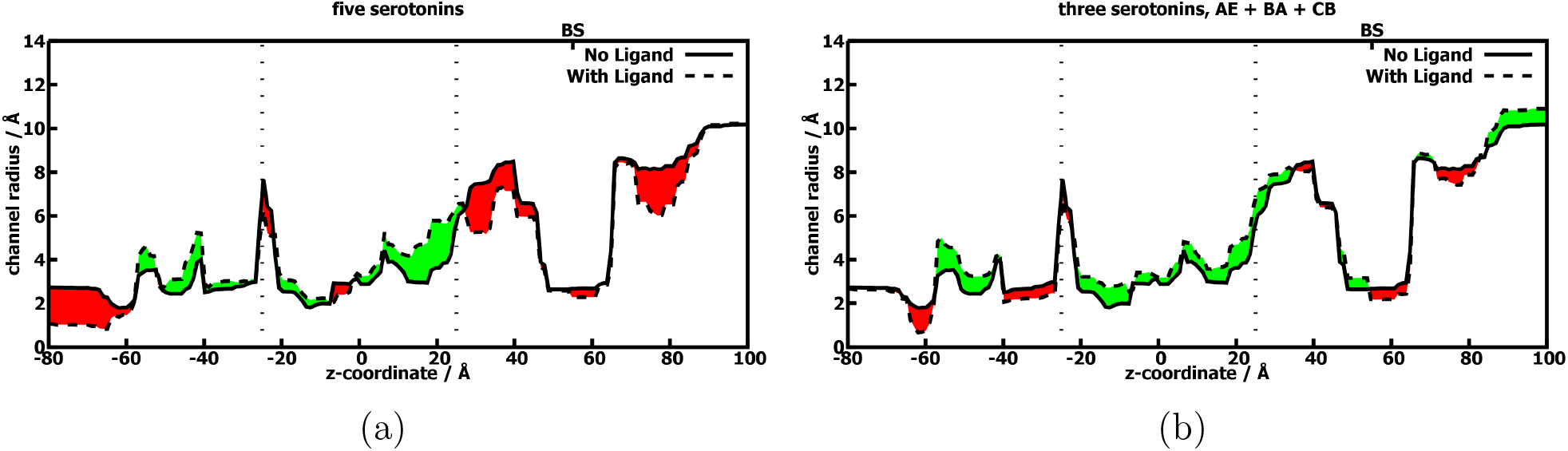
Diagram showing the predicted ion channel radius along the length of the 5-HT_3A_ receptor with (a) five and (b) three adjacent serotonin molecules bound. The solid line represents the data for the simulated equilibrium apo-structure, denoted by *μ*, the average position across all *N*_*t*_ snapshots. The broken line represents the profile for the receptor bound to three 5-HT molecules in adjacent binding sites. The figure legends are as in Figure 2. The five identical subunits of the 5-HT_3A_ receptor are denoted A to E in a clockwise direction when viewed from the extracellular space towards the cytoplasm. The ligands bind to the space between the subunits, and the binding site is denoted by the subunits adjacent to the binding site.

### 3.2 Antagonist binding

We applied our method for the competitive antagonists alosetron (PDB code: 6W1J), granisetron (PDB code: 6NP0), ondansetron (PDB code: 6W1M), and palonosetron (PDB code: 6W1Y). These four antagonists produced similar responses in the channel radius profile both experimentally and computationally, varying the number of antagonists bound also produced similar results (see Supplementary Figures).

Figure 4 shows the experimental results for the 5-HT_3A_ structure with five granisetrons bound (PDB code: 6NP0) in comparison to the experimental apo-structure (PDB code: 6BE1). The bottleneck radius in the transmembrane domain decreases from 2.63 Å to 2.55 Å, the rest of the transmembrane domain opens slightly but the binding of the competitive antagonists negligibly affects the ion channel. We evaluated the simulated structures in the same manner as we did the agonists. With five granisetron molecules bound, as shown in Figure 5, the channel bottleneck radius decreases from 1.81 Å to 1.67 Å. There is an increase in channel radius at −10 Å *< z <* 10 Å as well as at 30 Å *< z <* 40 Å in the extracellular domain.

**Figure 4.**
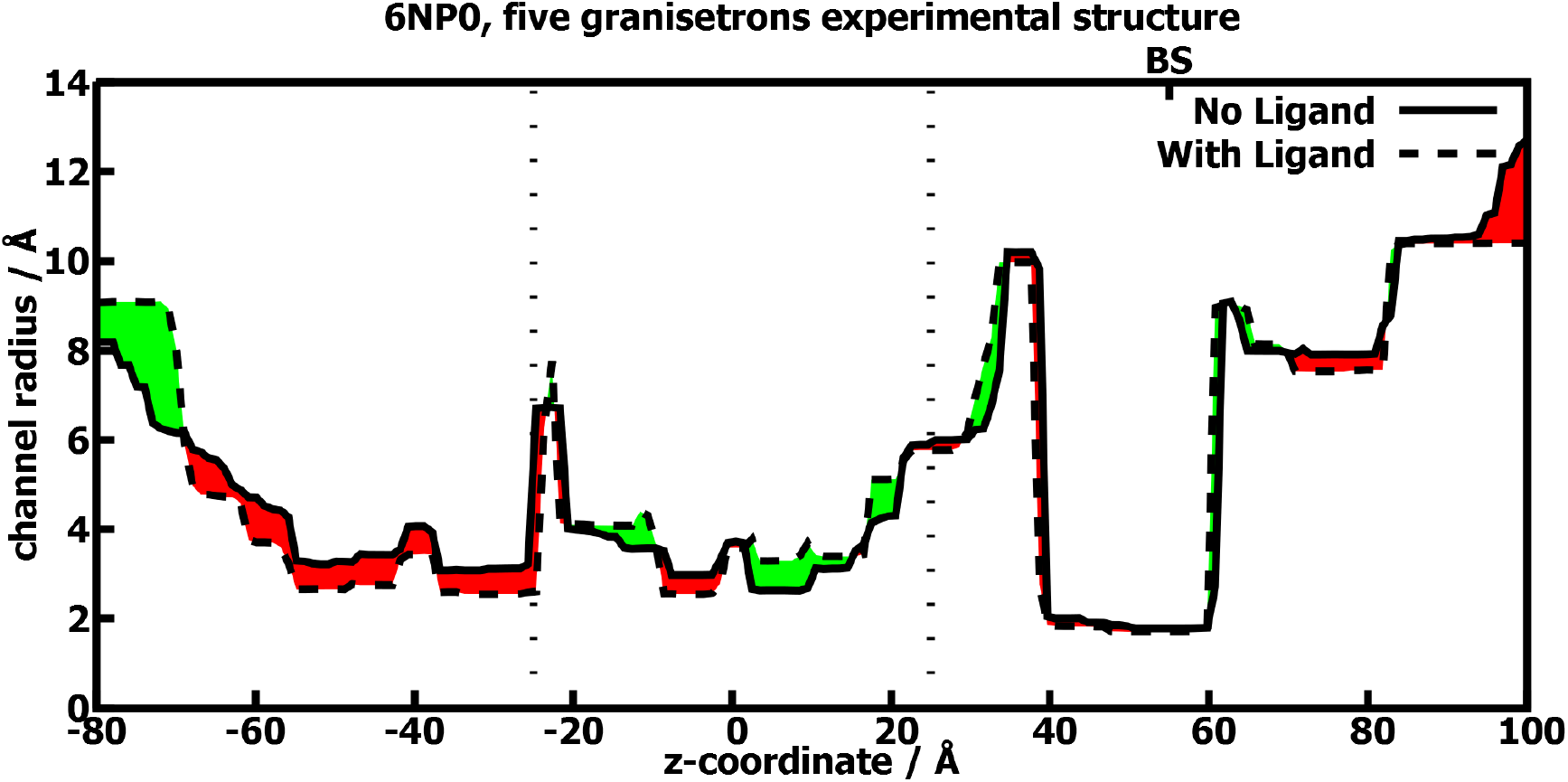
Diagram showing the radius of the ion channel along the length of the 5-HT_3A_ receptor. The solid line represents the data for the experimental apo-structure 6BE1. The broken line represents the profile for the receptor bound to five granisetron molecules 6NP0. The figure legends are as in Figure 2.

**Figure 5.**
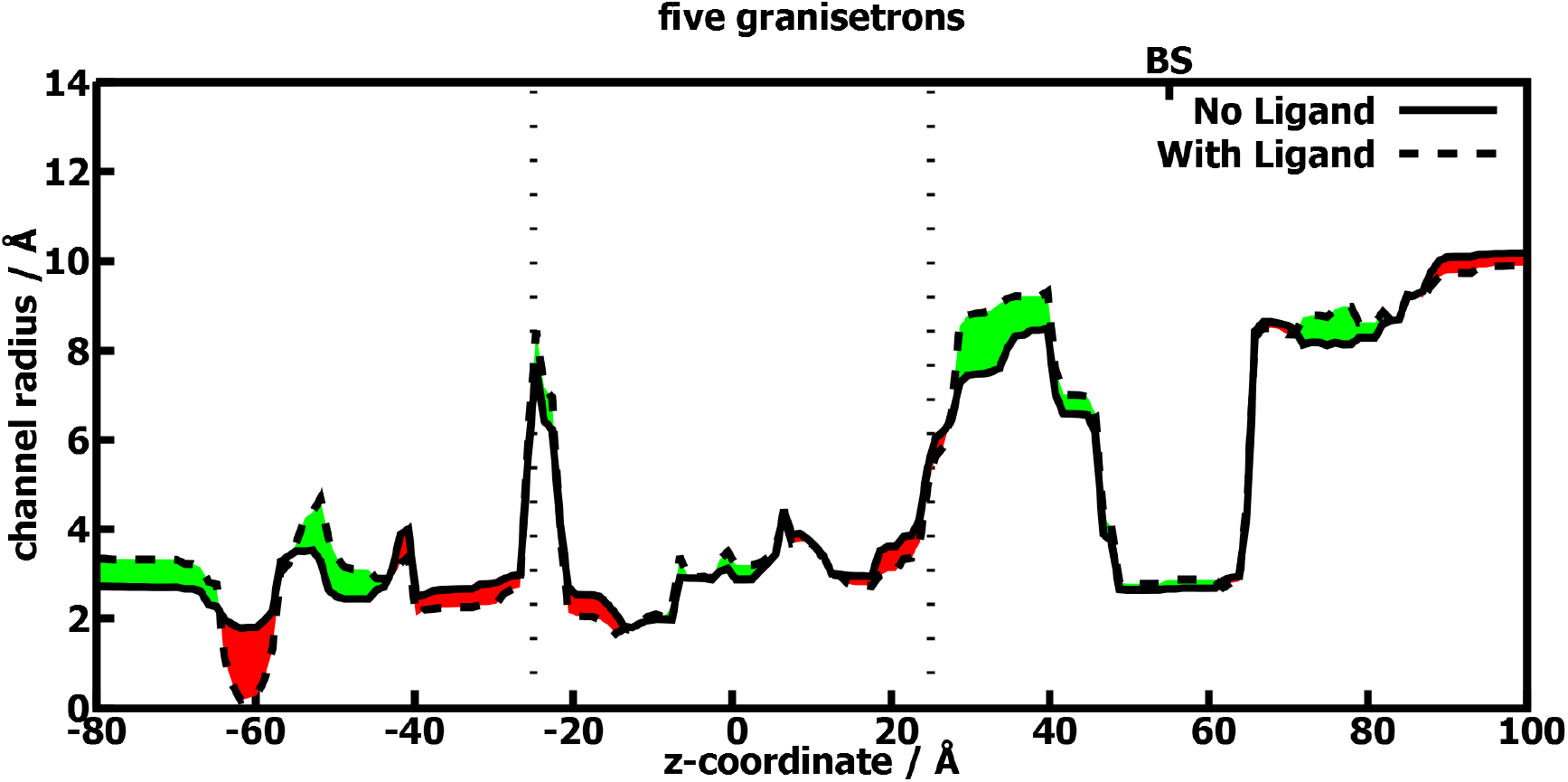
Diagram showing the radius of the predicted ion channel along the length of the 5-HT_3A_ receptor with five granisetron molecules bound. The solid line represents the data for the simulated equilibrium apo-structure *μ*. The broken line represents the profile for the receptor bound to five granisetron molecules. The figure legends are as in Figure 2.

## 4 Discussion

What is an agonist and what is a competitive antagonist? How do they act on the receptor to exert their effects? These are questions that have exercised the minds of pharmacologists for a long time. In this work, we have built upon a method to predict the gating effects (if any) of a ligand on its receptor. By gating, we mean the action of an agonist on a receptor which leads to the activation of its receptor. In this case, it is the binding of agonists which leads to the opening of the central ion channel.

Previously, structure determination experiments showed the effects of different ligands on the 5-HT_3A_ receptor and how different domains of the receptor were moved to either open the central ion channel, keep it closed, or even close it down further. ^5,9^ Further studies showed that this opening is probably asymmetric in the homopentamer.^6^

Maio et al. ^10^ used 4PIR as the starting structure, placed it in a hydrated membrane performed molecular dynamics simulations and free energy calculations. They showed that ion permeation was through the five lateral channels in the intracellular domain, the same region in which we see a sharp peak in the channel radius at *z* = −20 Å in the experimental structures. The experimental structure of the intracellular domain is not well resolved in the 5-HT_3A_ receptor, this combined with the lack of membrane in our simulations may explain the limitations of our method not detecting this.

Várnai et al. ^17^ used a model of the GABA_A_ receptor, carried out a CRANKITE simulation on the protein and developed a correlation tensor to relate agonist binding to gating; they did not study the effect of antagonists on the receptor. Our current work extends previous results^17^ by providing a fast and general method to study the effects of agonists and competitive antagonists on a receptor. As opposed to molecular dynamics studies, the use of CRANKITE does not require a large amount of computer time. The application of linear response theory is a general method: once we have calculated the covariance tensor, we can evaluate the perturbation caused by a ligand to the binding site, and thus predict the effect of the ligand; to study another ligand, we can use the same covariance tensor but re-evaluate the binding site perturbation. This makes it possible to rapidly define the effect of a large number of ligands on the same receptor. Moreover, we use the perturbed structure of the binding site rather than ligand induced forces as our input, which provides a better intuition for what novel drug candidates should look like. Lastly, our methods are able to accurately differentiate the behaviour of agonists and competitive antagonists: when agonists bind, there is widening of the ion channel in the transmembrane domain; and when competitive antagonists bind, there is no opening in the ion channel bottleneck.

The assumptions of our model only hold for weak perturbations in allosteric interactions. The perturbation must be weak enough, such that the higher order terms of the Taylor expanded potential are indeed negligible. The interaction must also take place sufficiently far away from the active site, so that there are no direct ligand-induced forces acting on the site of interest.

In the future, it would be interesting to develop a method to reverse the question: if one would like to open an ion channel, in what ways should one perturb the binding site? If one were to develop a competitive antagonist, what freedoms does one have to perturb the binding site without causing activation?

## Supporting information

Predicted channel radius profiles of the 5-HT3A receptor after applying our method to a variety of ligands, along with experimental profiles.

## 5 Acknowledgements

The authors thank Patrick Welche, Michele Simoncelli, and Clara Wanjura for their physical insight, Nigel Unwin and Ian Martin for useful discussion pertaining to receptor biology, and Stuart Rankin and Michael Rutter for help with computing. Simulations in this work were carried out using resources provided by the Cambridge Service for Data Driven Discovery (CSD3) operated by the University of Cambridge Research Computing Service, provided by Dell EMC and Intel using Tier-2 funding from the Engineering and Physical Sciences Research Council (capital grant EP/P020259/1), and DiRAC funding from the Science and Technology Facilities Council. W.J.J. acknowledges funding from the Rod Smallwood Studentship at Trinity College, Cambridge.

## 7 Supplementary Material

Supplementary Figures: Predicted channel radius profiles of the 5-HT_3A_ receptor after applying our method to a variety of ligands, along with experimental profiles.

