## Supplementary material for "Differentiating Agonists and Competitive Antagonists of the Serotonin Type 3A (5-HT_3A_) Receptor": Predicted channel radius profiles of the 5-HT3A receptor after applying our method to a variety of ligands, along with experimental profiles.

### 1 Supplementary Figures

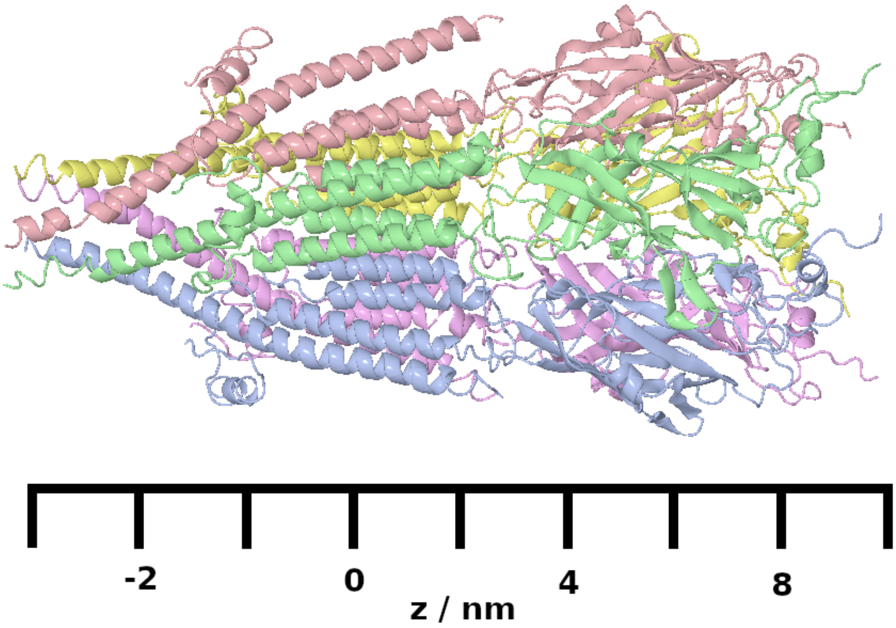


Figure 1: Diagram showing the 5-$HT_{3A}$ receptor with a coordinate system aligned along the receptor's long axis. The extracellular domain is in the region $z > 25$ Å, the transmembrane domain is in the region $-25$ Å$\leq z \leq$25 Å and the intracellular domain is in the region $z < -25$ Å. The agonist and competitive antagonist binding site is situated between individual subunits and is in the region $z = 50$ Å.

### 1.1 Experimental Channel Profiles

Diagrams show the radius of the ion channel along the length of the 5-$HT_{3A}$ receptor. The solid line represents the data for the experimental apo-structure 6BE1. The broken line represents the profile for the receptor bound to five ligands. The transmembrane domain, in the region $-25\,\text{Å} < z < 25\,\text{Å}$, lies within the broken vertical lines. BS denotes the position of the binding site. The spaces between the solid and broken lines are colored red for a reduction in the ion channel radius or green for an increase in the ion channel radius.

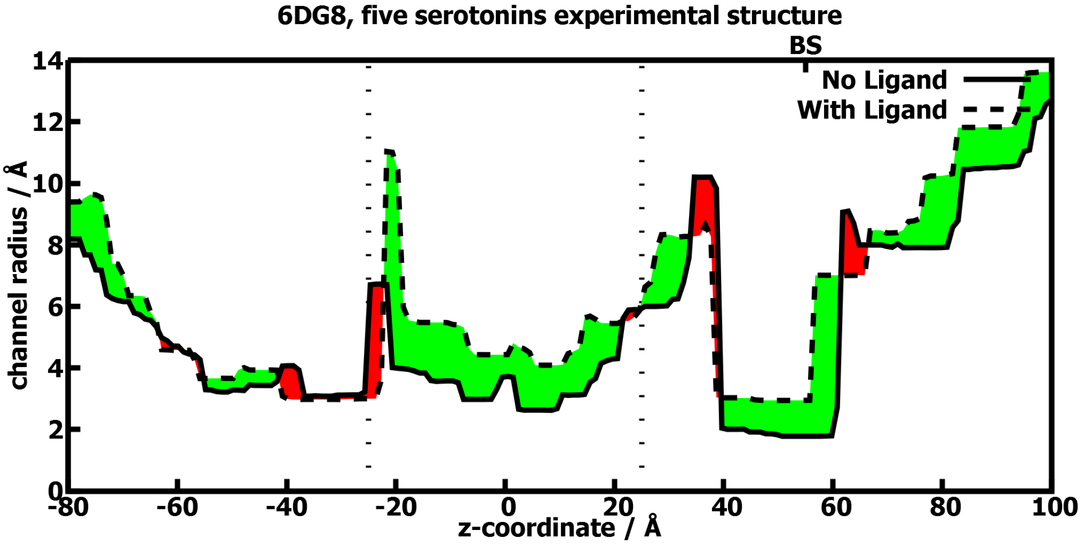


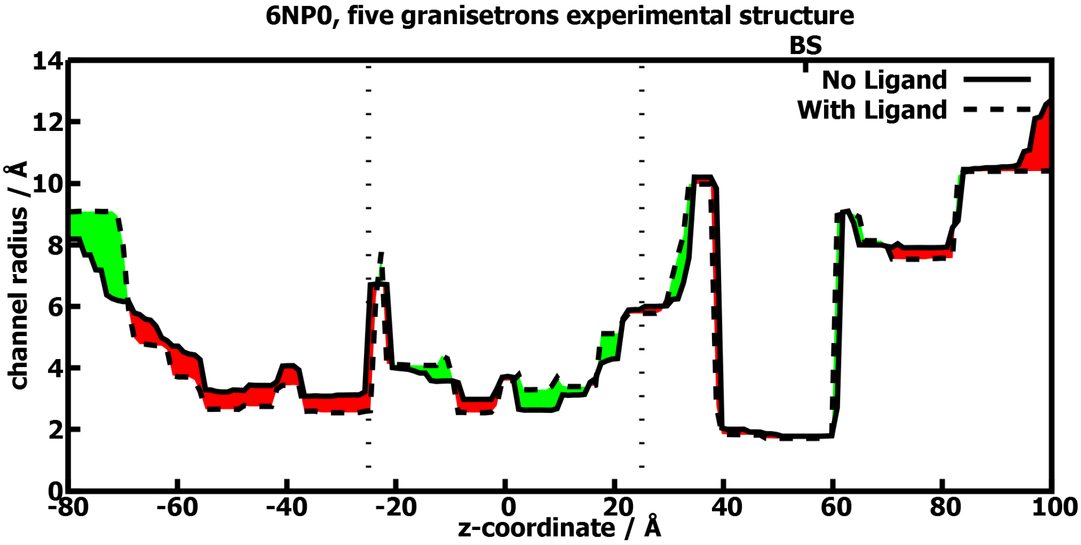

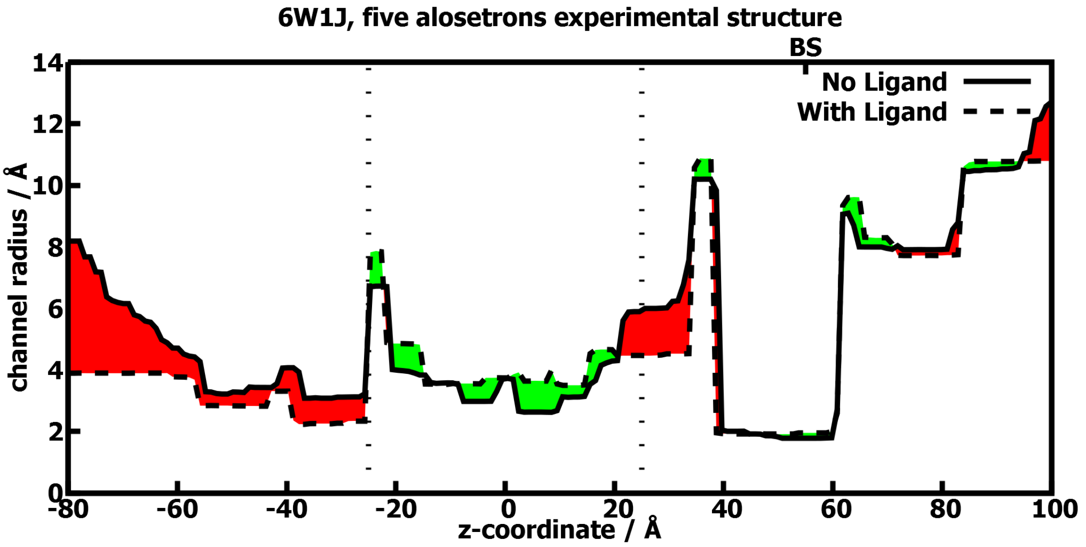
6W1J, five alosetrons experimental structure
BS
No Ligand
With Ligand
channel radius / Å
z-coordinate / Å

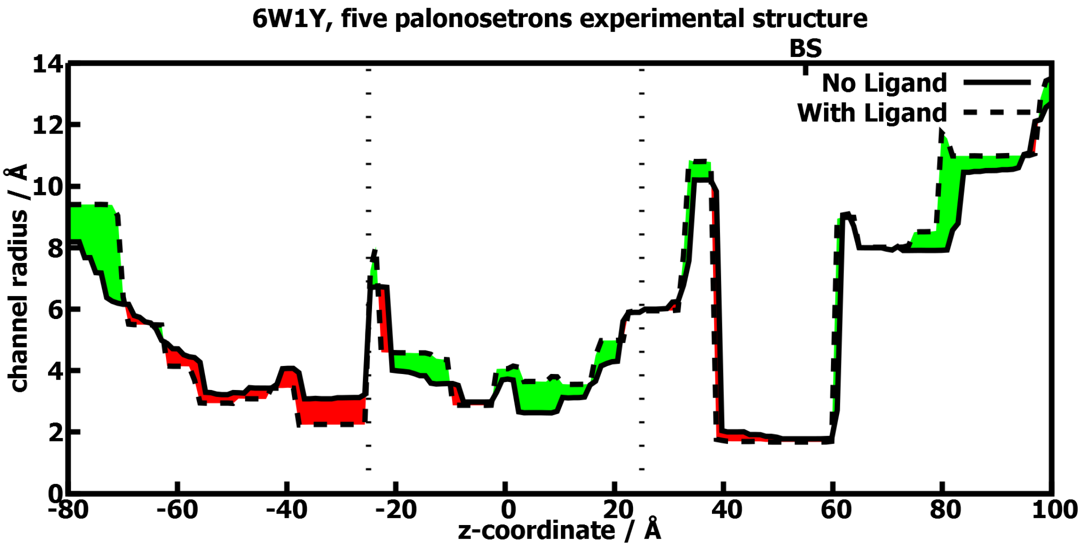
6W1Y, five palonosetrons experimental structure
BS
No Ligand
With Ligand
channel radius / Å
z-coordinate / Å

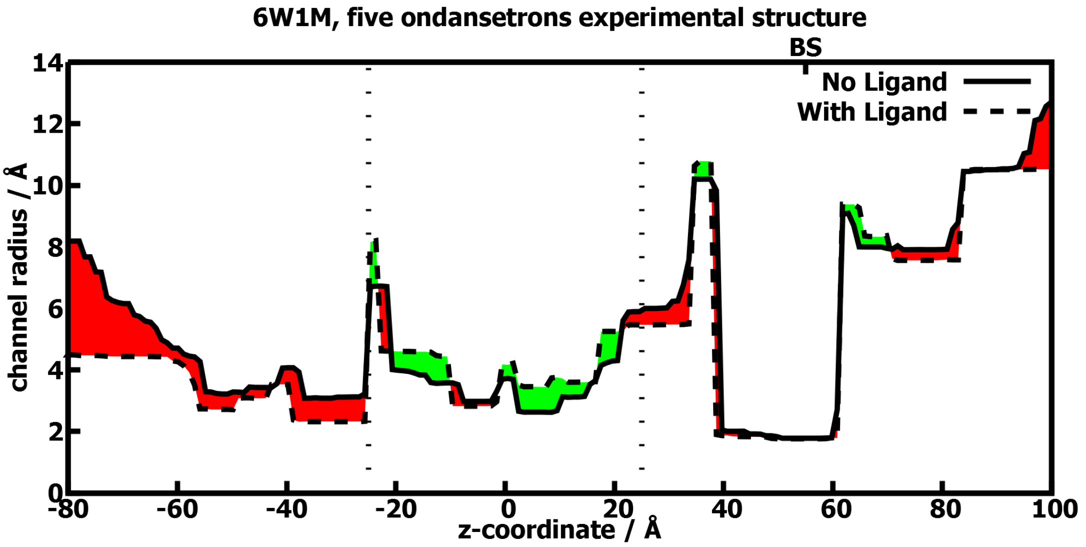
6W1M, five ondansetrons experimental structure
BS
No Ligand
With Ligand
channel radius / Å
z-coordinate / Å
0
2
4
6
8
10
12
14
-80
-60
-40
-20
0
20
40
60
80
100

### 1.2 Predicted Channel Profiles

Diagrams show the radius of the ion channel along the length of the 5-$HT_{3A}$ receptor. The solid line represents the data for the simulated equilibrium apo-structure, taken as the ensemble average, $\mu$. The broken line represents the profile for the receptor bound to the specified ligand(s). The five identical subunits of the 5-$HT_{3A}$ receptor are denoted A to E in a clockwise direction when viewed from the extracellular space towards the cytoplasm. The ligands bind to the space between the subunits, and the binding site is denoted by the subunits adjacent to the binding site.

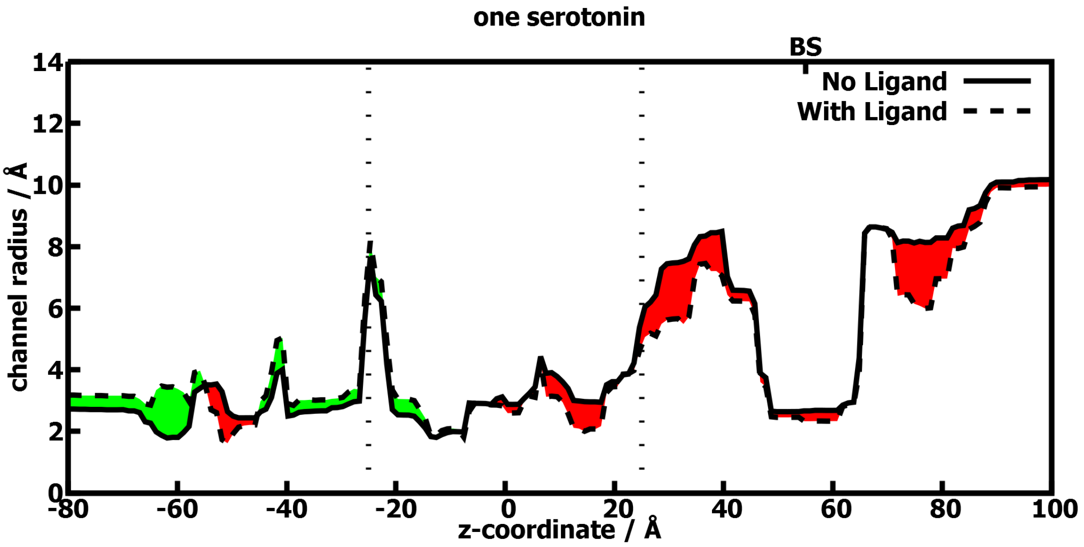


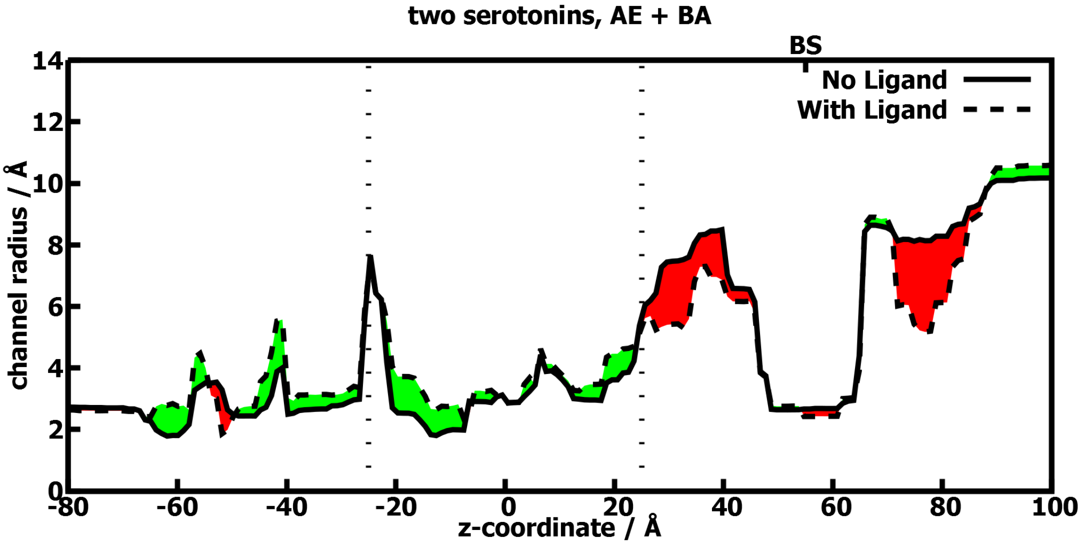

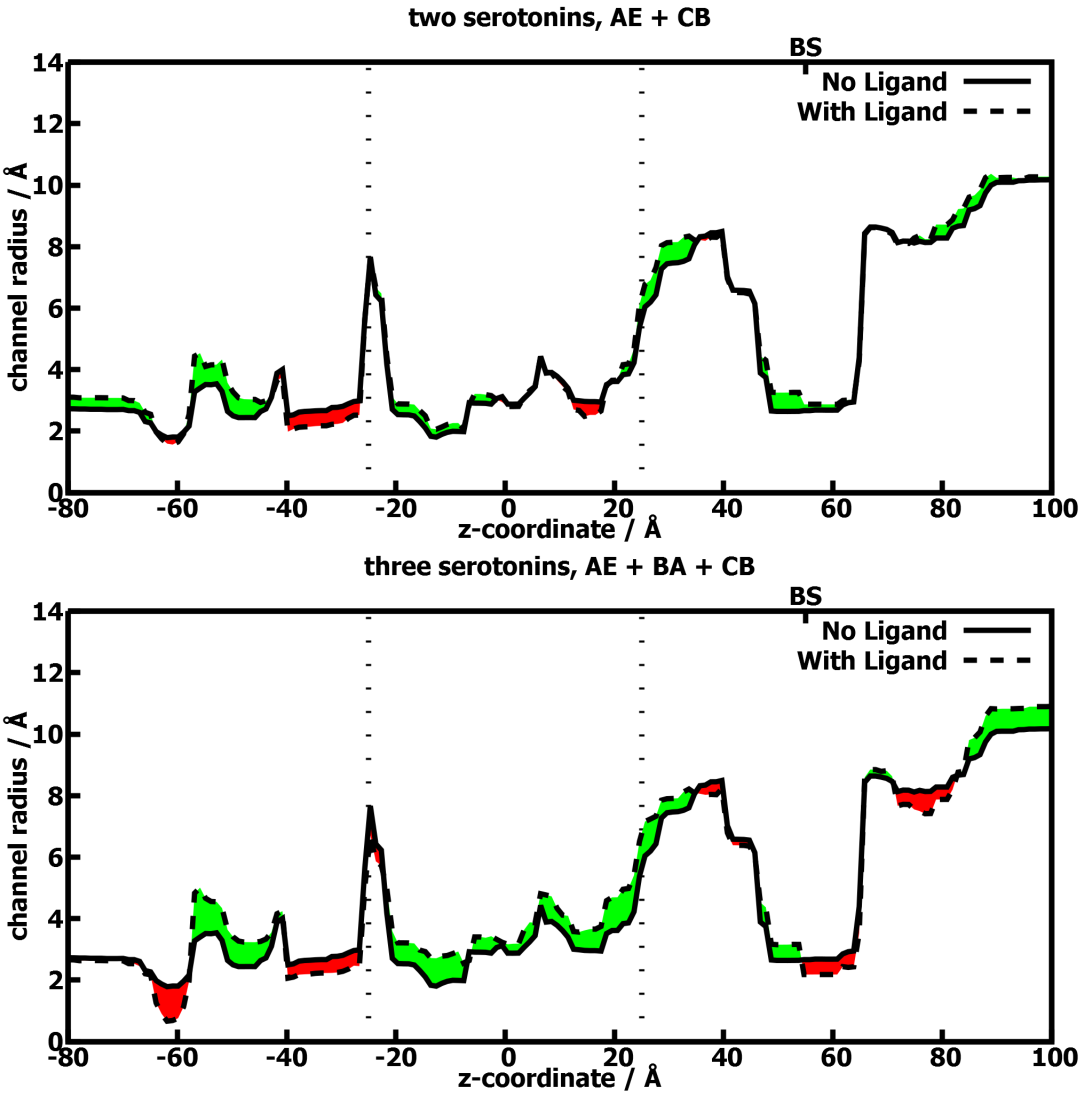

two serotonins, AE + CB
BS
No Ligand
With Ligand
channel radius / Å
z-coordinate / Å
0
2
4
6
8
10
12
14
-80
-60
-40
-20
0
20
40
60
80
100
three serotonins, AE + BA + CB
BS
No Ligand
With Ligand
channel radius / Å
z-coordinate / Å

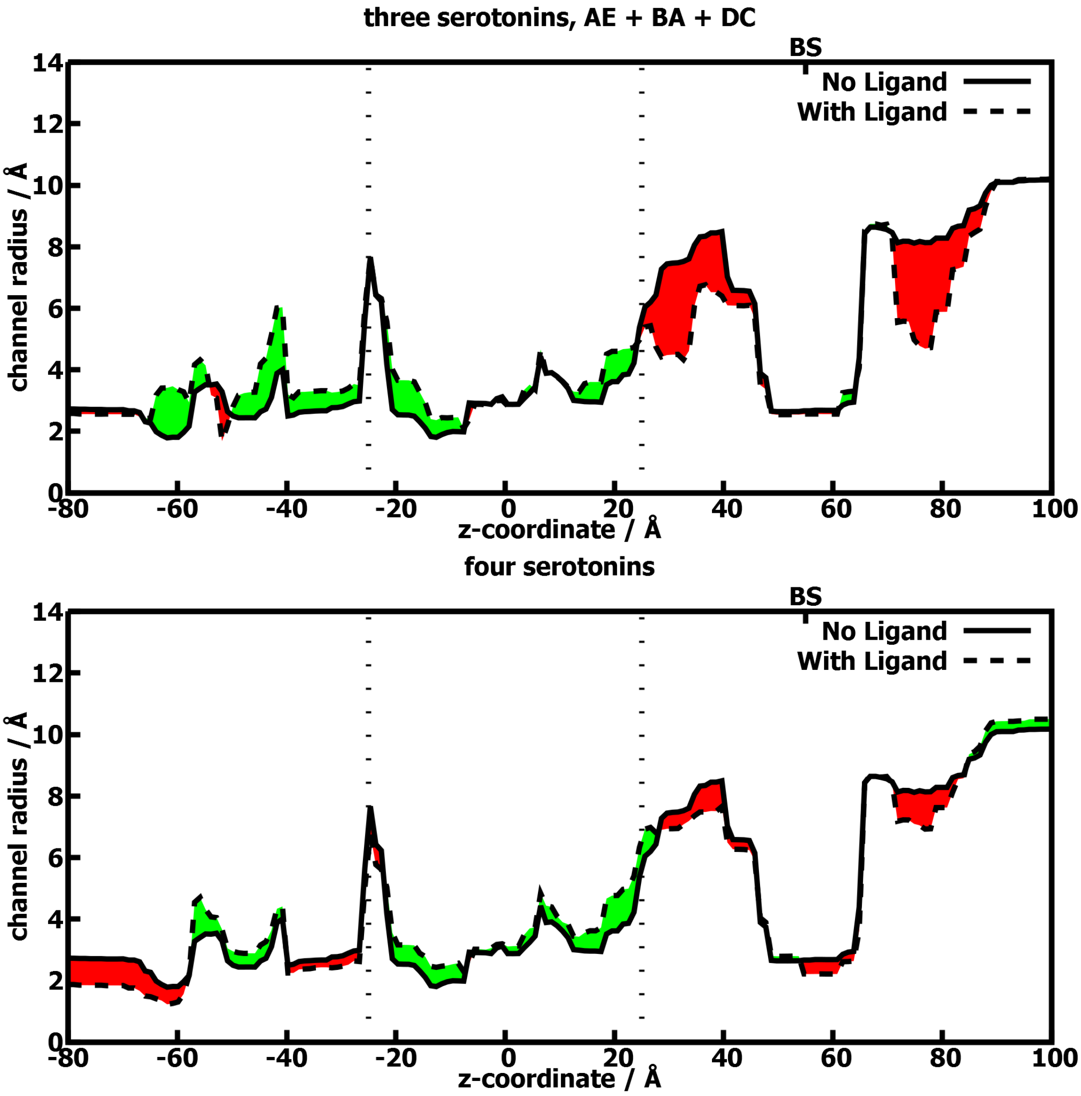
three serotonins, AE + BA + DC
BS
No Ligand
With Ligand
channel radius / Å
z-coordinate / Å
four serotonins
BS
No Ligand
With Ligand
channel radius / Å
z-coordinate / Å

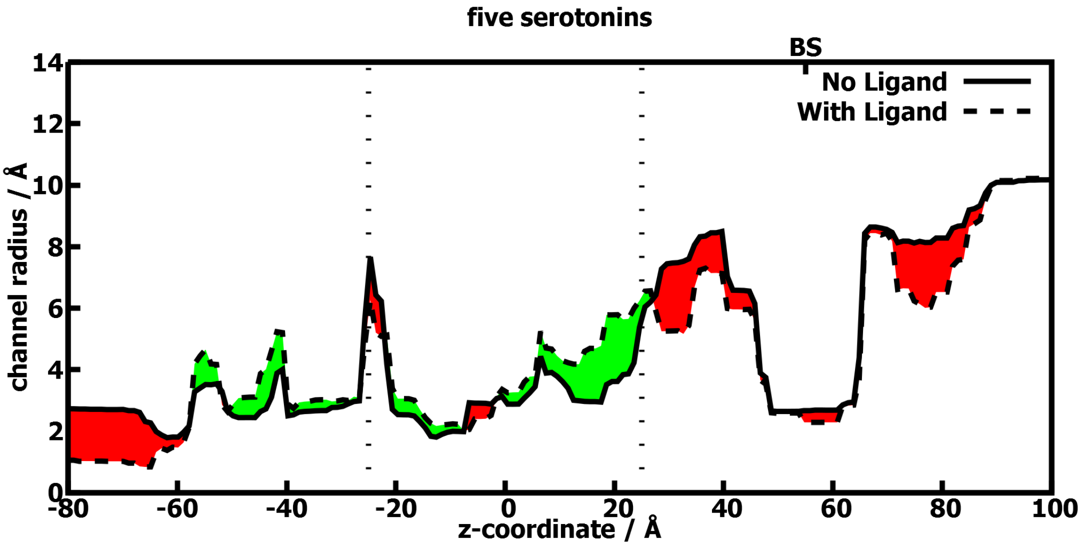
five serotonins
BS
No Ligand
With Ligand
channel radius / Å
z-coordinate / Å
0
2
4
6
8
10
12
14
-80
-60
-40
-20
0
20
40
60
80
100

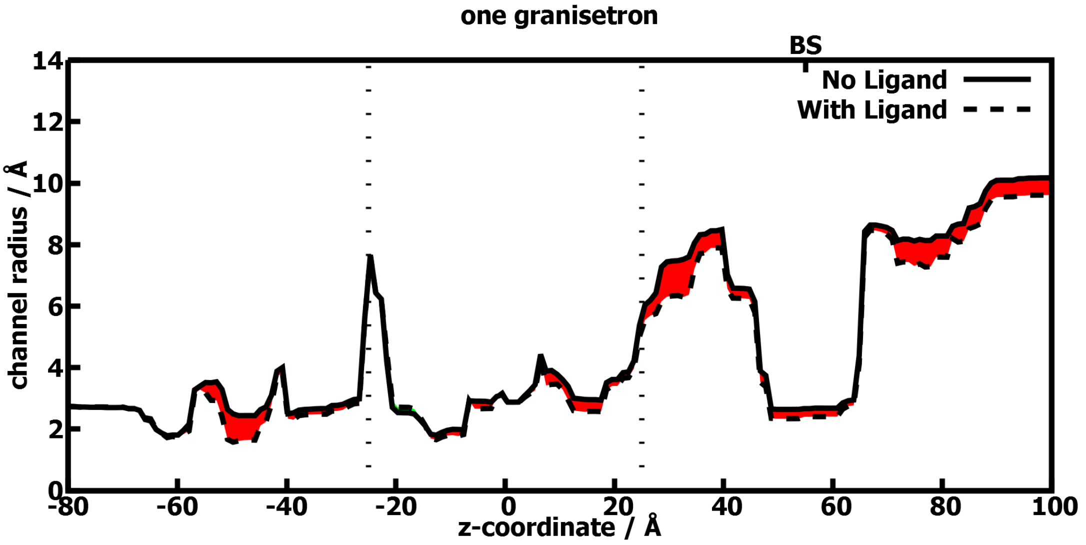
one granisetron
BS
No Ligand
With Ligand
channel radius / Å
z-coordinate / Å
0
2
4
6
8
10
12
14
-80
-60
-40
-20
0
20
40
60
80
100

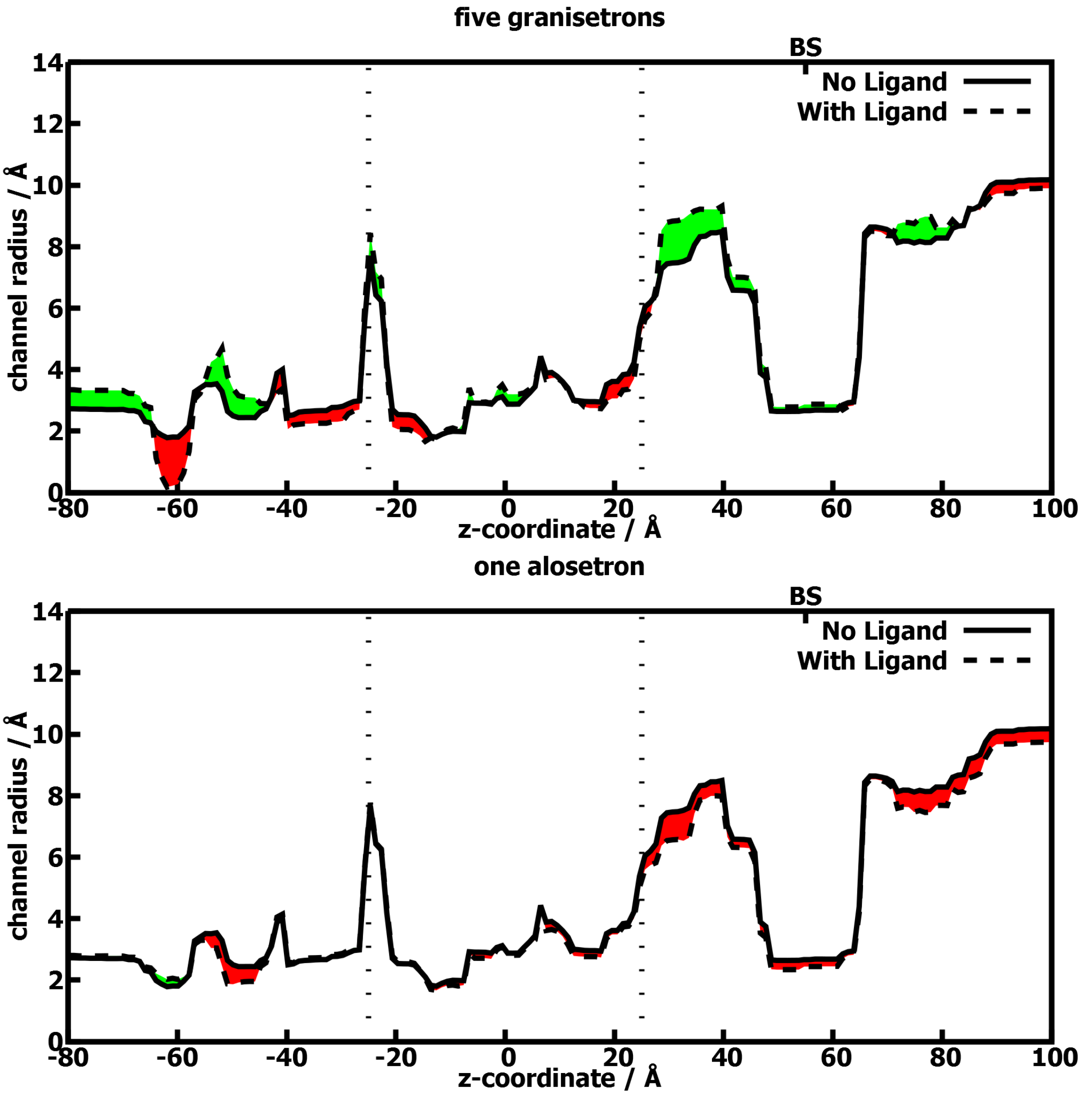
five granisetrons
BS
No Ligand
With Ligand
channel radius / Å
z-coordinate / Å
one alosetron
BS
No Ligand
With Ligand
channel radius / Å
z-coordinate / Å

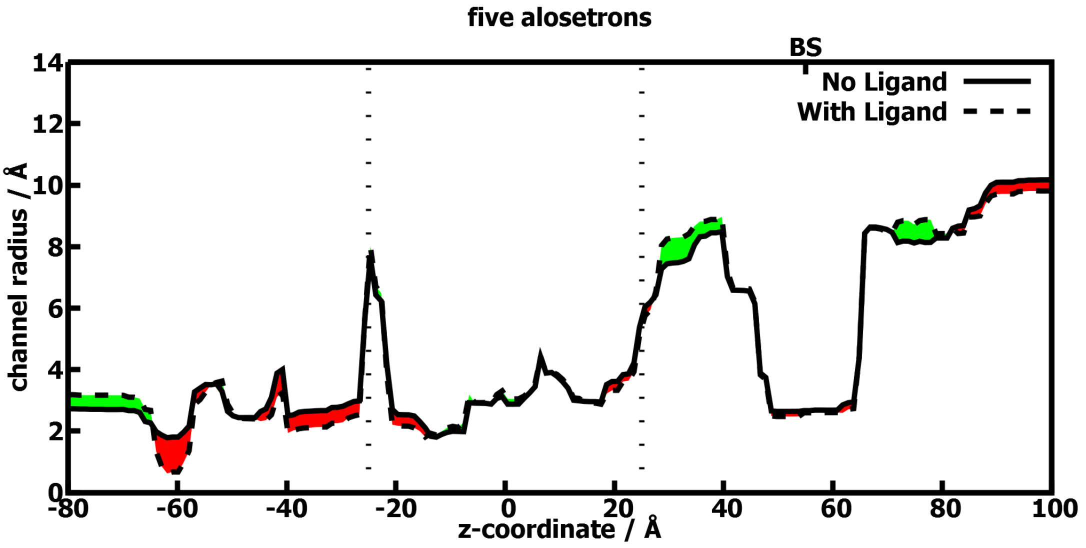
five alosetrons
BS
No Ligand
With Ligand
channel radius / Å
z-coordinate / Å
0
2
4
6
8
10
12
14
-80
-60
-40
-20
0
20
40
60
80
100

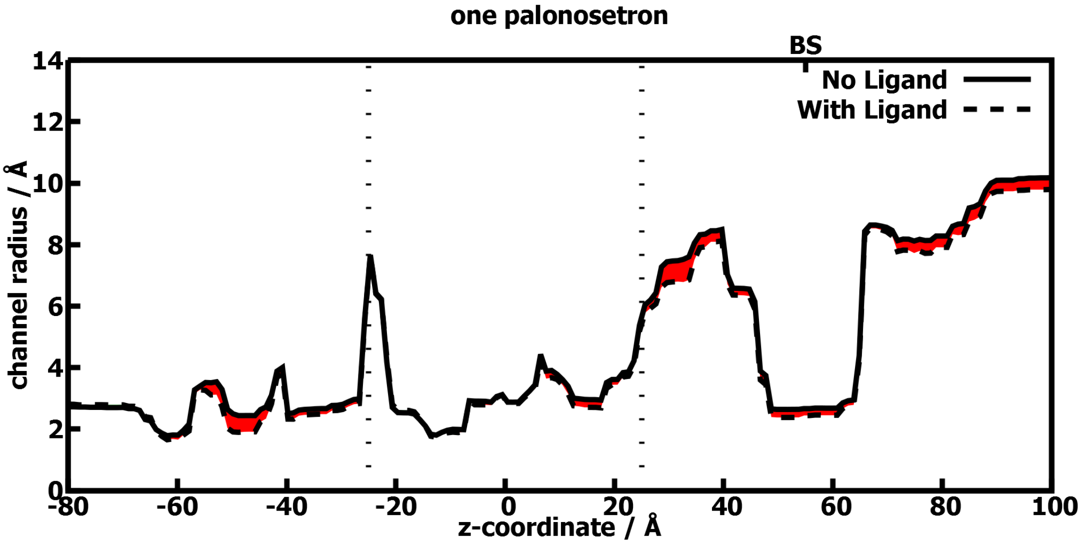
one palonosetron
BS
No Ligand
With Ligand
channel radius / Å
z-coordinate / Å
0
2
4
6
8
10
12
14
-80
-60
-40
-20
0
20
40
60
80
100

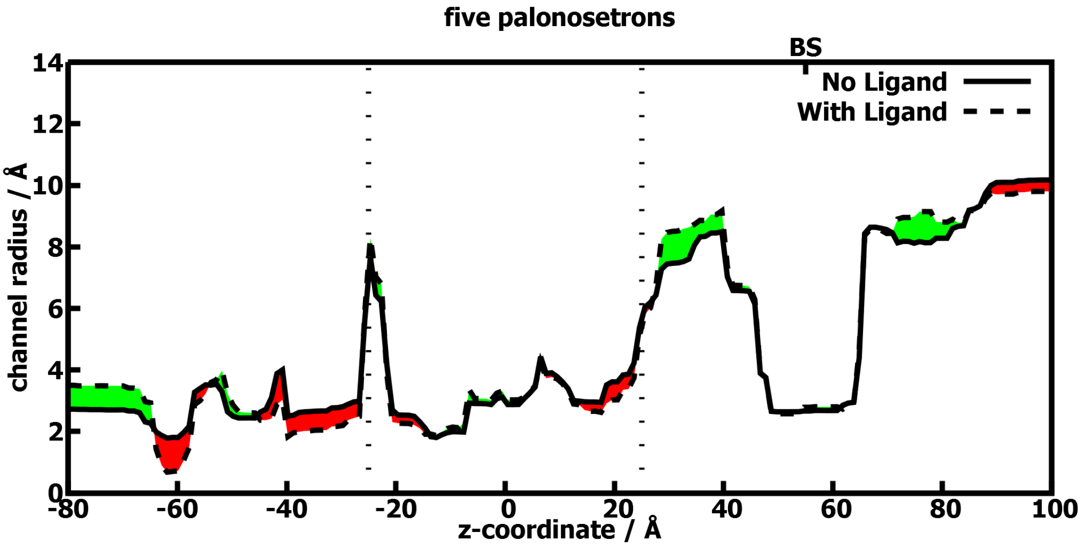
five palonosetrons
BS
No Ligand
With Ligand
channel radius / Å
z-coordinate / Å
0
2
4
6
8
10
12
14
-80
-60
-40
-20
0
20
40
60
80
100

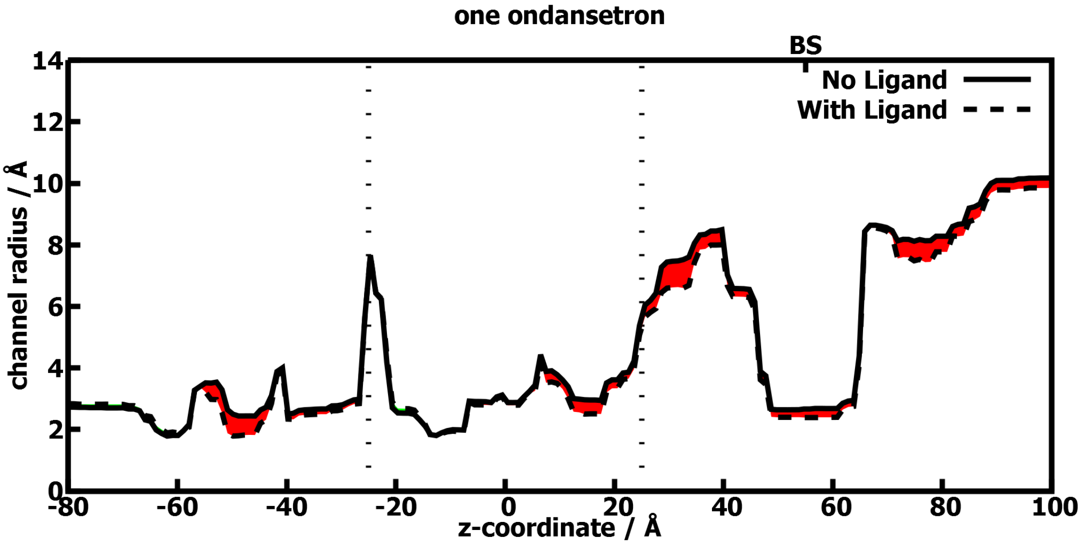
one ondansetron
BS
No Ligand
With Ligand
channel radius / Å
z-coordinate / Å
0
2
4
6
8
10
12
14
-80
-60
-40
-20
0
20
40
60
80
100

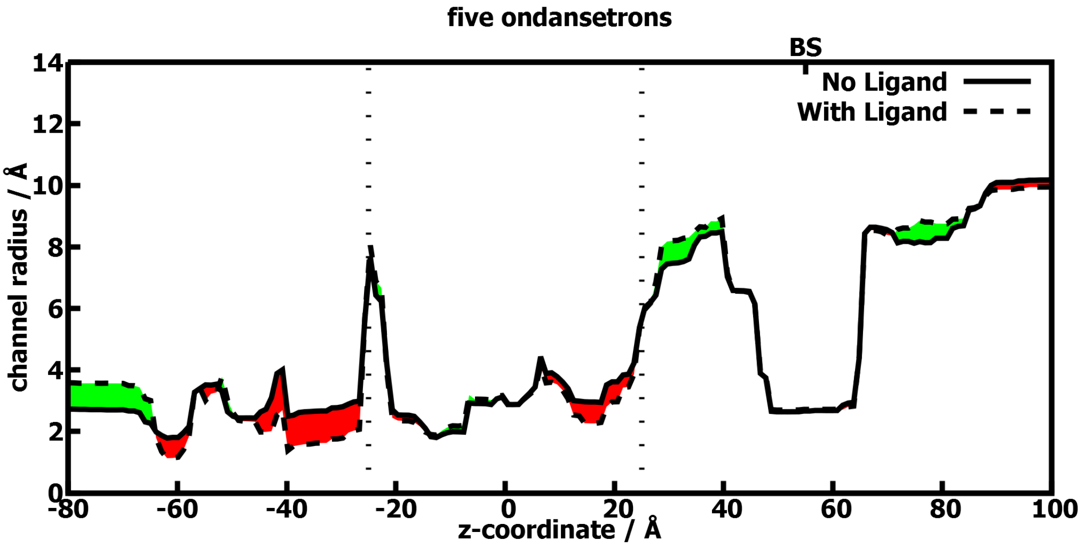
five ondansetrons
BS
No Ligand
With Ligand
channel radius / Å
z-coordinate / Å
0
2
4
6
8
10
12
14
-80
-60
-40
-20
0
20
40
60
80
100
